# Scientific reasoning driven by influential data: resuscitate *dfstat*!

**DOI:** 10.1101/2024.10.30.621016

**Authors:** Andrej-Nikolai Spiess, Stefan Rödiger, Matthias Schaks, Michał Burdukiewicz, Joel Tellinghuisen

## Abstract

In biomedical literature, one of the most widely employed statistical procedures to analyze and visualize the association between two variables is linear regression. Data points that exert influence on the fit and its parameters are routinely, but not as often as required, identified by established influence measures and their corresponding cut-off values. In this work, we are specifically concerned with the presence of influential data points that directly impact hypothesis testing of linear regressions, which none of the established measures describe. Interestingly, the highly overlooked influence measure *dfstat* and its derived leave-one-out *p*-value exists exactly for this purpose, unmentioned in the majority of statistical text books as well as absent from all available statistical software packages. Its application for identifying these data points seems pivotal, as scientific reasoning in publications is almost exclusively based on the *p*-value of the fit, commonly adhering to the α = 0.05 threshold to state significance or not. With this metric, we found for 29 of 100 digitizable papers published in *Science, Nature* and *PNAS* in 2016, a time when the “reproducibility crisis” was a growing concern, that stated significances (or their absence) are based on the presence of a single influential data point.

## Introduction

In the 2010’s, increasing discussion on the existence of a “reproducibility crisis” emerged, sparked by failing to replicate 70 % of published preclinical studies in two commercial wet-lab efforts (Begley et al., 2012; Prinz et al., 2011), lack to reproduce results from 10 out of 18 microarray studies (Ioannidis et al., 2009), non-confirmation of several psychological/sociological studies through actual experiments (Shanks et al., 2015), estimating a reproducibility of 0.6 to 6.8 % for nearly 2000 hydrology papers (Stagge et al., 2019), as well as statistical reanalyses uncovering discrepancies between original and replicated effect sizes/*p*-values (Open Science Collaboration, 2015; Camerer et al., 2018). As this crisis was widely percepted in the scientific community, it was unsurprising that in a survey of 1576 researchers in *Nature*, 90 % of scientists confirmed a reproducibility crisis, with own or other’s experiments not repeatable in 50-90 % of cases (Baker, 2016).

A frequent procedure in research is to test the association between two variables through linear regression. We noticed – from simple “eyeballing” – a surprising number of biomedical research articles in high impact journals where the visual display of the regression analysis suggested the presence of influential data points potentially affecting the regression outcome, specifically in terms of significance statements. Influential data points usually deviate substantially from the main data cloud, strongly affect the regression outcome and can, in the optimal case, be identified by influence measures such as *hat values* (leverage), *dffits* (change in fitted values), *dfbeta(s)* (change in parameter estimates), *covratio* (change in parameters’ covariance matrix), *Cook’s distance* (changes in leverage and residuals) and *studentized* residuals (Cook, 1977; Cook & Weisberg, 1982). Each of these measures is associated with certain cut-off values, determined from the measure’s value, sample size and number of parameters (Belsley et al., 2004), however caution is advised (Fox, 1991).

In biomedical research, the association’s significance is often the primary statement, in which case suspected influential data would ideally be examined for their effect on the regression’s *p*-value. This translates into the question: How precisely does the slope’s *p*-value change with and without the questionable data point(s)? Here, the null hypothesis,

H_0_: *β*_*1*_ = 0 (no association) is tested with the statistic 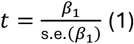 with *n*-2 degrees of freedom, where *β*_*1*_ and s.e.(*β*_*1*_) are the slope and its standard error, respectively. The common threshold for significance is α = 0.05, ubiquitously employed for making the dichotomous statement of significant association or not, but the wisdom of this choice has been questioned (McShane et al. 2017; Nuzzo, 2014).

In this work, using the neglected influence measure *dfstat* (Belsley et al., 1980) and its derived evaluation of leave-one-out *p*-values, we aimed at providing a quantitative assessment on the frequency of linear regressions depicted in high impact journals in the year 2016 whose statistical conclusions were sensitive to single data points. Specifically, from a body of 66, 87 and 100 publications of year 2016 mentioning “linear regression” in the journals *Science, Nature* and *PNAS*, respectively, we reanalyzed those graphs displaying well-separated and digitizable data points for outliers that are influential on hypothesis testing.

## Results & Discussion

Our literature search of the journals *Science, Nature* and PNAS of year 2016 resulted in 24/66, 30/87 and 46/100 linear regression graphs, respectively, which were analyzable through complete and digitizable data point separation. Undoubtedly, our graph/data selection entailed substantial bias to lower sample sizes as they not only appear, but actually are, more amenable to digitization through more discernible and less overlapping data points. As the original raw data values were not supplied except in two publications, the online graphs were exported as tiff-files, digitized using a graph digitizer (WebPlotDigitizer) and raw values exported (Supplementary Data 1; “Science Data”, “Nature Data” and “PNAS Data”). This approach was consequential from the experience that biomedical raw data is rarely supplied in the original articles (Iqbal et al., 2016; Miyakawa, 2020) or frequently held back on request to the authors (Allison et al., 2016; Wicherts et al., 2011). The following applies to all 100 digitized graphs, however in Supplemental Data 1 we have, for conciseness, noted those cases where influential data was present: each dataset was subjected to an initial linear regression with all data points to confirm that the obtained regression parameters (sample size, *p*-value, Pearson’s *r*, Spearman’s *ρ*, coefficient of determination *R*^*2*^) matched closely to those stated in the original paper (Supplementary Data 1, “Results”, “Stated in publication” *versus* “Acquired from digitization”). Here, some small discrepancies were found with respect to sample size and Spearman’s *ρ* (Supplementary Data 1, “Results”, “Comments”), and it was conspicuous that for some publications, essential regression parameters such as sample size, *R*^2^, exact *p*-values or effect sizes (slope) were not provided (cells with asterisks in Supplementary Data 1, “Results”, “Stated in publication”). The *p*-values derived from the digitized linear regressions deviated from those stated in the publications in the third decimal place (*e*.*g*., 0.08/0.086; 0.02/0.023; 0.03/0.036).

We then applied *dfstat* to the digitized data, in order to examine the effect of single data points on hypothesis testing. Evidently, *dfstat* is living in oblivion: it is not common knowledge, even in the statistical community, that there exists an influence measure pertaining directly to the change in *t*-statistics, that will “show whether the conclusions of hypothesis testing would be affected” (Belsley et al., 1980), termed *dfstat* (Belsley et al., 1980; Baltagi, 2011; Temple, 2000) or *dfstud* (Rousseeuw & Leroy, 1987):

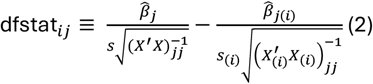

where 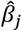 is the *j*-th estimate, *s* is the residual standard error, *X* is the design matrix and (*i*) denotes the *i*-th observation deleted. *dfstat*, which for an univariate regression’s slope *β*_1_ is the difference of *t*-statistics 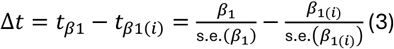, is therefore linked to leave-one-out changes in *p*-value, Δ*p*, calculated from

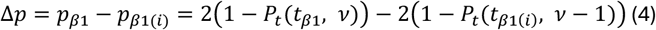

where *Pt* is the Student’s *t* cumulative distribution function with *v* degrees of freedom. When the α-threshold is used for dichotomizing into “significant” or “non-significant”, a change in statistical conclusions occurs when either *p*_*β*1_ <*α*< *p*_*β*1*(i)*_ or *p*_*β*1*(i)*_ <*α*< *p*_*β*1_.

In applying this procedure to all points *i*, similar to the established influence measures, we identify data points influential on the significance conclusion, by paying attention to the crossing of the α*-*threshold. Figure 1 shows such an analysis for the digitized data from Figure 1 in Paper-01. Deletion of the data point at 10/3.89 (orange point) shifts the original *p*-value from 0.043 (green line) to 0.17 (orange line). Furthermore, four classical influence measures – *dfbetas, dffits, covratio* and *studentized residuals* – exceed their commonly used cut-off values (Belsley et al., 1980) for this data point and it is outside the 95 % prediction interval of the *n* -1 model which excludes it (Figure 1, dashed lines).

**Fig 1.**
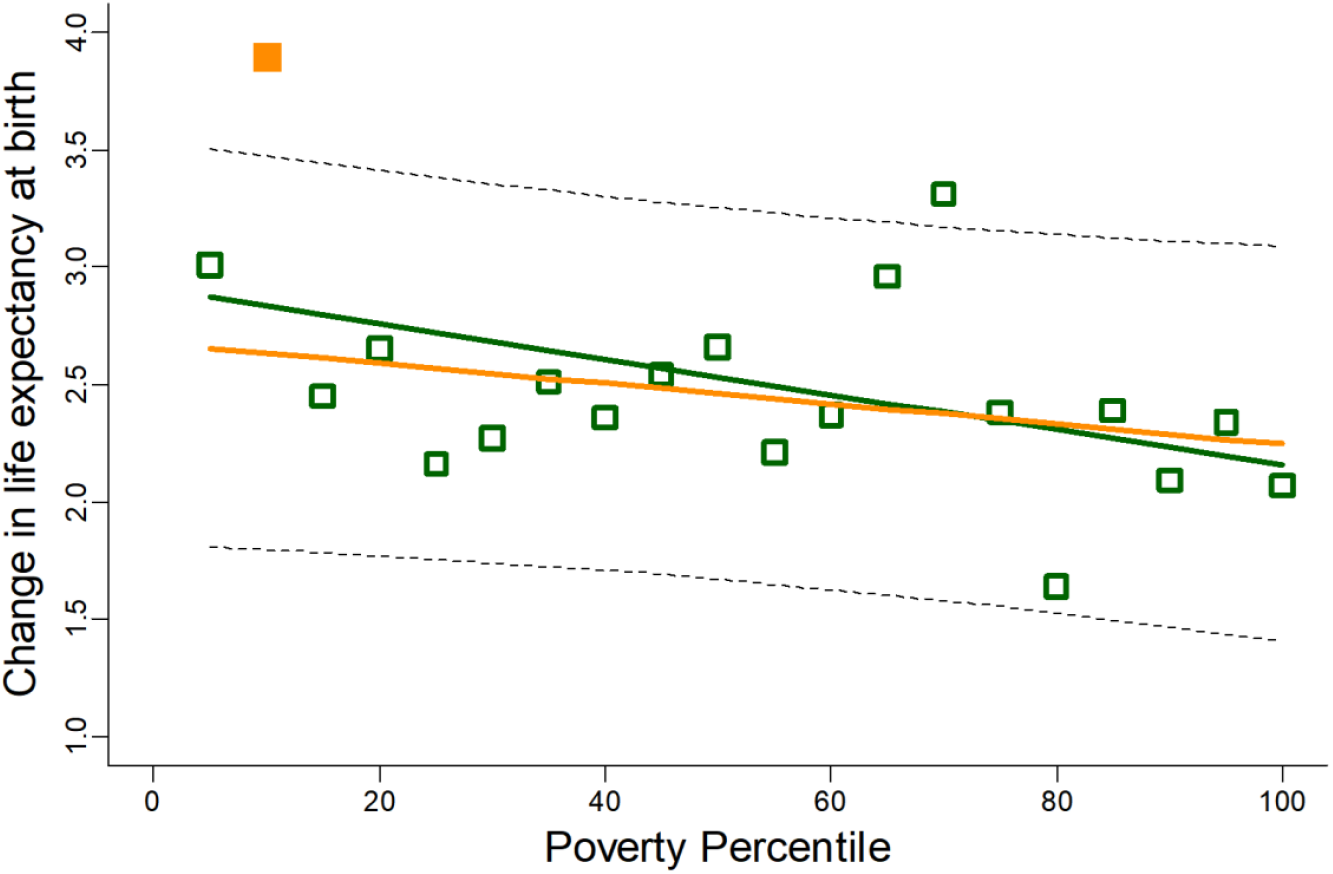
Digitized version of Figure 1 in Paper-01 with influential data point marked in orange. Green and orange trend lines correspond to regression with (*p* = 0.043) and without (*p* = 0.17) influential data point, respectively. Dashed lines: 95 % prediction interval.

The above finding is surprisingly frequent: for the year 2016 and the 100 publications in *Nature, Science* and PNAS that we inspected, we found 21 publications (29 graphs), for which the same characteristics applied. When omitting their most influential data point with respect to changes in *p*-value (large Δ*p* with effect on significance statement), we could often observe drastic changes (Figure 2; difference between 2nd (green) and 3rd (orange) bar), emphasizing that the original significance and conclusions derived thereof were merely supported by the presence of a single influential data point (!). Specifically, a change from significant to insignificant was encountered in 18 cases, and *vice versa* for the other 11 cases.

**Fig 2.**
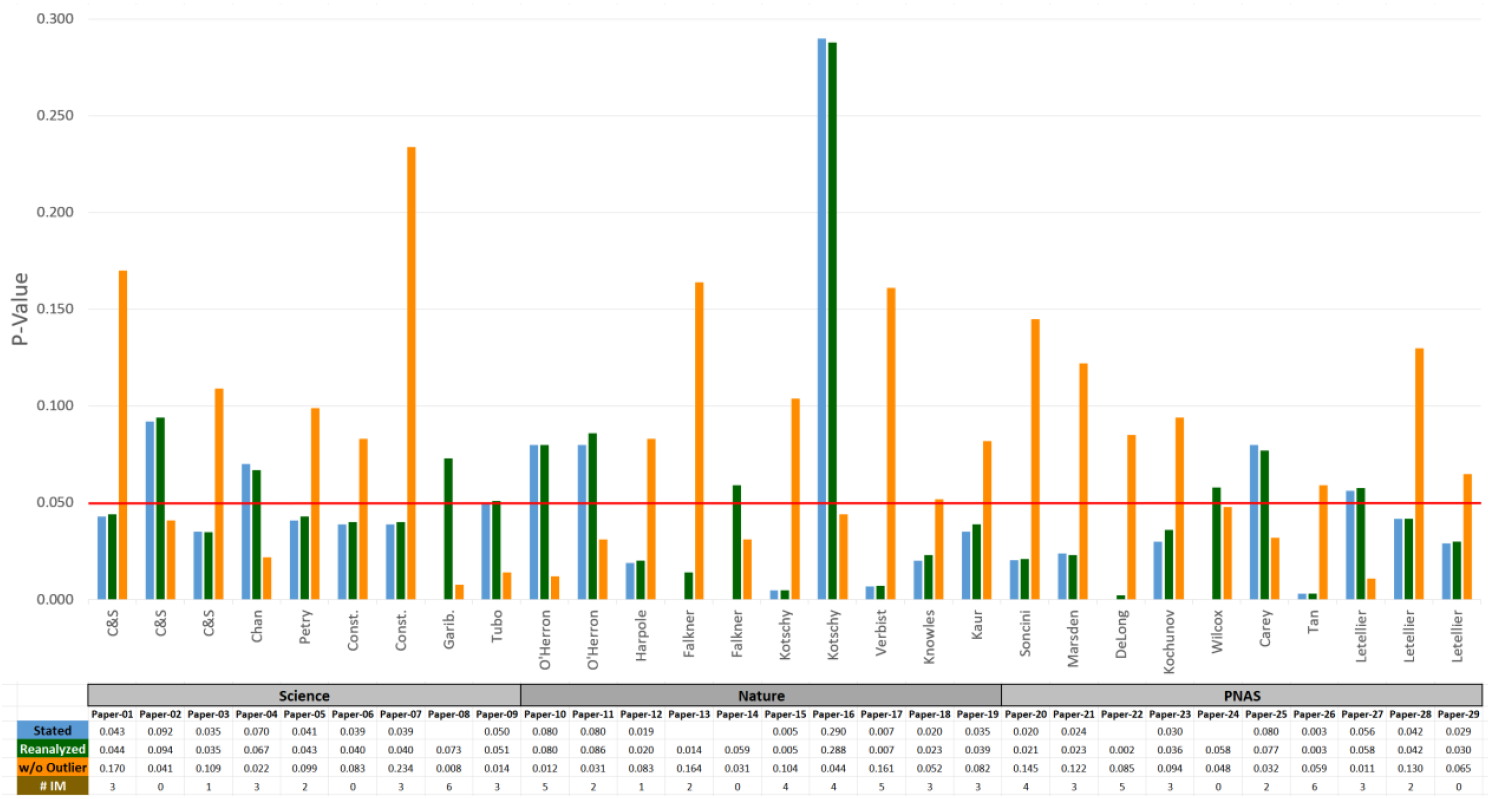
Upper panel: *P-*values stated in the original publication (blue), reanalyzed after digitization (green) and after removal of the highest influential data point (orange). Lower panel: *P*-values of the above panel in tabular format; # IM: number of influence measures exceeding their threshold. The red horizontal line demarcates α = 0.05.

The relatively high incidence of these changes in our re-analyzed publications (∼ 20 %) strongly advocates for a mandatory similar treatment in linear regression-derived significance statements. Interestingly, *dfbeta*(slope) exceeded its specific cut-off value of 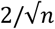 (Cook, 1977) in 21 of 29 cases for the most influential data point. For 25 in 29 cases (86.2 %), such data points would have been identified by one and often more of the six classical influence measures in combination with their established cut-off values (Figure 2, lower panel, “# IM”), indicating a considerable overlap between classical influential data points and those that alter the significance statement upon deletion. None of the reanalyzed publications mentions checking their data in this manner, thus we conclude that influential data analysis is largely neglected in this field.

Studies on the effect of influential data points on the regression’s *p*-value are limited. Hollenbeck *et al*. (2006) focused on the *p*-values of Pearson’s correlation *r*, to which similar conclusions in this work apply through the known identity of the *t*-statistics for *r* and *β*1 (for derivations see Pugh, 1966; Soch, 2024). Choi (2009) demonstrated the effect of omitting *dfbeta*-identified outliers on the slope and *p*-values of foreign direct investment-related variables. Unfortunately, none of the common statistical software packages (*e*.*g*., SPSS, SAS, Stata, Matlab, R, Python) implements a leave-one-out influence analysis on the regression’s *p*-value.

For more insight on how *dfstat* and its derived leave-one-out *p*-value compares to the classical influence measures, we simulated three different regimes where we added single data points into a 100 x 100 grid surrounding regression data, thereby mimicking influential data points with differing severity. In the first regime, we created a *null* model (*n* = 20; *R*^2^ = 0, *p* = 1) and consecutively added values within a 100 x 100 grid of *x*_i_ ∈ [-50, 50] and *y*_i_ ∈ [-1, 1] as influential data points. A subsequent analysis of *dfbeta*(slope), *dffits, covratio, hat value*, Cook’s distance and leave-one-out *p*-value shows characteristic patterns for each of the influence measures (Figure 3), either when the measure’s cut-off value is exceeded (Figure 3A-E, orange areas) or when the regression’s *p*-value is rendered significant (Figure 3F, green area). The latter green areas with *p* < 0.05 reveal that for the classical influence measures, the same corresponding areas would exhibit values exceeding their cut-off, corroborating our findings with respect to excessive dfbeta(slope) values when the significance statement is changed.

**Fig 3.**
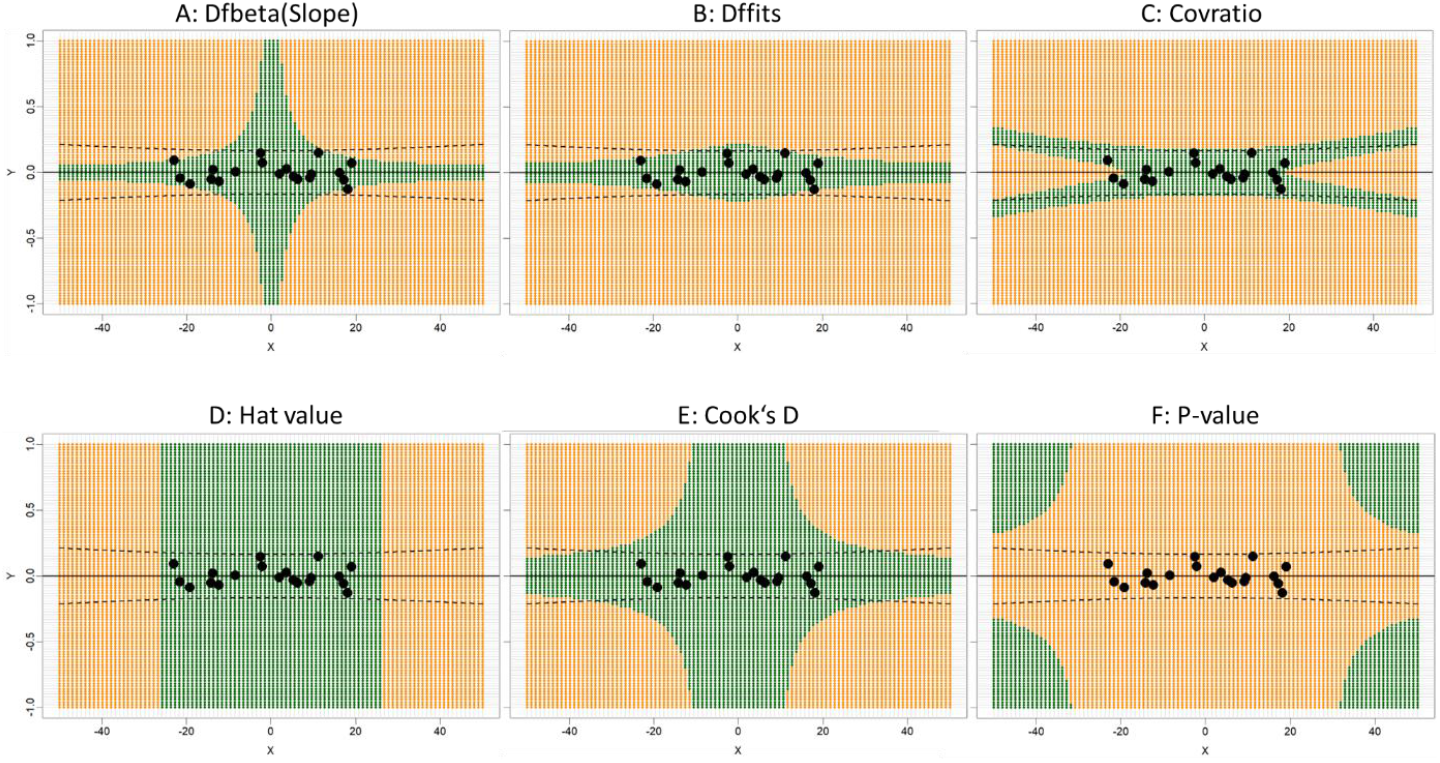
Analysis of the effect of influential data points from a 100 x 100 grid surrounding a null model on the values of dfbeta(slope) (A), dffits (B), covratio (C), hat value (D), Cook’s D (E) and regression’s *p*-value (F). Data points in orange areas would result in exceeding the corresponding cut-off value (A-E), those in green in rendering the regression’s *p*-value significant (F). Dashed lines: 95 % prediction interval.

In the second regime (*n* = 20), we inspected the effect of influential data points when the regression’s *p*-value is approaching the α-threshold (Supplemental Figure 1). Here, we can observe that the area of attaining significance is increasing when the *p*-value is in proximity of the α-threshold (Supplemental Figure 1, A-D, green area) and intersecting with a higher proportion of the 95 % prediction interval (dashed lines). Conversely, when the regression is already significant (Supplemental Figure 1, E-F), decreasing *p*-values result in a narrowing of the area where an influential data point might render the regression insignificant (orange). Essentially, these findings are along the lines of the unstable *replication probability* (Boos et al., 2011; Miller, 2009) or “fickle” *p*-values (Halsey et al., 2015) in proximity of the α-threshold. It also tallies with the simulations in Supplemental Figure 1 and the relatively high frequency of *p*-values near the α-threshold encountered in our reanalyzed studies (Figure 2) and in general biomedical literature (Masicampo, 2012). Furthermore, an increasing proximity to the α-threshold entails an increasing number of data points that are rendered influential, as demonstrated by simulating a wide range of *p*-values for *n* = 20, 200 (Supplemental Figure 2, A+B).

In the third regime, based on an insignificant model (*p* = 0.1), we evaluated the effect of increasing sample sizes (*n* = 10, 20, 50, 100, 200, 500) on the susceptibility of the model to be altered with respect to the significance statement through an influential data point. As somewhat expected, with increasing sample size, the area for influential data is migrating further away from the data cloud (Supplemental Figure 3 A-F, green area), meaning that it takes either higher leverage on the independent variable *x* or higher studentized residuals on the dependent *y* value to change the significant statement. Note that also here, at small sample sizes (*n* = 10, 20), the area of influential data is partly within the data cloud and intersects with a considerate proportion of the 95 % *n* - 1 prediction interval (dashed lines).

To this end, we feel compelled to give the following summary:

i) in the realm of classical influence measures, there is none which describes the effect of influential data point on hypothesis testing.
ii) *dfstat* and its associated leave-one-out *p*-value are neither routinely employed as influence measures in univariate regression nor in Pearson correlation analysis.
iii) cases where single data points drive the significance statement are not rare, even in high-end journals.
iv) there is an extensive overlap in location where a data point exceeds classical influence measure cut-offs and where a change in significance statement occurs.
v) resembling considerations from *replication probability*, a proximity of the *p*-value to the α-threshold entails less distance of an influential data point to the main data cloud to exert an effect on the significance statement.
vi) with increasing sample size, the effects of influential points on changes in *p*-values progressively diminish.

The primary problem we see in the overlooked identification of influential data points that drive the significant statement is that they form the basis for unambiguous conclusions, such as “…, indicating increasing inequality in life expectancy over this period” (Currie & Schwandt, 2016; *p* = 0.043)” or “… suggesting that resource addition caused changes in community composition that were not always associated with diversity loss” (Harpole et al., 2016; *p* = 0.019). Therefore, when α = 0.05 is set as a strict decision boundary in light of unstable data, and statistical significance is used as the “coin of the realm” in many areas of investigation, this poses the risk of directing future research on the presence of single influential data points that exert regression significance. Conversely − and at least equally important − is that influential data-driven insignificant regressions may not be published or omitted from further inspection. Ways to meliorate this problem have been suggested by a number of scientists, such as i) put less emphasis on *p*-values (Wasserstein & Lazar, 2016; Blakeley et al., 2019), ii) be less strict with dichotomizing thresholds (Betensky, 2019; Nuzzo 2014), iii) identify, mark and mention influential data points (Altman & Krzywinski, 2016) or iv) substitute/supplement least-squares regression with robust alternatives (Rousseeuw & Leroy, 1987).

With this work, we want to advocate the use of an influential measure that describes the effect of outliers on the significance statement. As long as stating significance or not is still based on the ubiquitous α = 0.05 threshold, these statements can be sensitive to the presence of a single data point, are highly unstable and most probably render the analysis less reproducible.

## Materials & Methods

### Literature Search

The term “linear regression” was used to browse the 2016 database of *Science, Nature* and PNAS, yielding 66, 87 and 255 results, respectively, with the following search terms (last accessed 2024-09-16): https://www.science.org/action/doSearch?field1=AllField&text1=linear+regression&ConceptID=&publication%5B%5D=science&publication=&Ppub=&AfterMonth=1&AfterYear=2016&BeforeMonth=12&BeforeYear=2016 https://www.nature.com/search?q=“Linear%20Regression”&journal=nature&date_range=2016-2016&order=relevance https://www.pnas.org/action/doSearch?field1=AllField&text1=“Linear+Regression”&field2=AllField&text2=&publication%5B%5D=pnas&publication=&Ppub=&AfterYear=2016&BeforeYear=2016&access=on

Specifically, the number of hits with the search term “linear regression” (“Hits”), number of graphs that were not shown (“Not shown”), number of graphs that were not analyzable (“Not analyzable”), number of graphs that were analyzable (“Analyzable”), number of Articles in which the analyzable graphs were found (“Articles”) and number of graphs with influential variables (“Influential variable”) were as given in Table 1.

**Table 1.** Characteristics from the literature search with the term “linear regression” and graphs that were re-analyzable.

|  | Hits | Not shown | Not analyzable | Analyzable | Articles | Influential variable |
| --- | --- | --- | --- | --- | --- | --- |
| **Science** | 66 | 23 | 19 | 24 | 6 | 9 |
| **Nature** | 87 | 19 | 38 | 30 | 7 | 10 |
| **PNAS** | 100 | 24 | 30 | 46 | 8 | 10 |

For PNAS, only the first 100 results were used for the following analysis. The graph analysis was limited to those with a visual impression of “digitizability”, meaning that only those were selected where the data points exhibited clear separation and no complete overlap.

### Digitization

Graph digitization was performed with the WebPlotDigitizer online software (see References),using the original online graph images (.jpg, .tif or .png format) from the publications. Attention was paid to accurate digitization by zooming into axis borders and the data points before marking them with the digitization tool. After digitization, the raw (*x, y*) data were exported as a text-file.

### Analysis of influence measures

The digitized raw data were imported in R (www.r-project.org) and analyzed with the `lmInfl` function of our reverseR package (see References. The console output then directly indicates whether an influential data point changes the significance statement when present or not, *e*.*g*.:

~~~
> res <-lmInfl(LM)
Model is significant at p = 0.007036032.
Found the following significance reversers, in order of
strength:
Point #11 => p = 0.1605117.
~~~

The function exports all classical influence measures and in addition *dfstat*, leave-one-out *p*-values, Hadi’s measure (Hadi 1992), Pena’s measure (Pena 2005) and the *coefficient of determination ratio* (Zakaria et al., 2014). Measures that exceed their specific thresholds are marked with asterisks. Results from these analyses, including the raw data from digitization, are to be found in Supplementary Data 1, Tabs “Science Data”/”Nature Data”/”PNAS Data”. The summary of the complete analysis including study names, analyzed figures, statistics as mentioned in the paper as well as after digitization, difference in stated and acquired *p*-value, statistics after removal of the highest influencer, direction of significance change and comments can be found in Supplementary Data 1, Tab “Results”.

### Simulations

Three different regimes were simulated in which we added single data points into a 100 x 100 grid surrounding regression data in order to mimick influential data points:

1) *Null* model from uniform distribution (𝒰_[-26, 20],_ *n* = 20; *R*^2^ = 4.3E-34, *p* = 1), iterating values in a 100 x 100 grid of *x*i ∈ [-50, 50] and *y*i ∈ [-1, 1] as influential data points.
2) non-*Null* models from uniform distribution (𝒰_[-26, 20],_ *n* = 20) with different *p*-values, iterating values in a 100 x 100 grid of *x*i ∈ [-100, 100] and *y*i ∈ [-5, 5] as influential data points.
3) non-*Null* models from uniform distribution (𝒰_[-26, 20],_ *p* = 0.1) with different sample sizes *n* = 10, 20, 50, 100, 200, 500, and iterating values in a 100 x 100 grid of *x*i ∈ [-100, 100] and *y*i ∈ [-5, 5] as influential data points.

We chose to employ uniform distributions for our simulations to obtain a dense point cloud. The choice of distribution is of marginal importance as the Gauss-Markov theorem for obtaining the *best least unbiased estimator* (BLUE) only puts emphasis on the distribution of residuals.

---

**Supplemental Figure 1.**
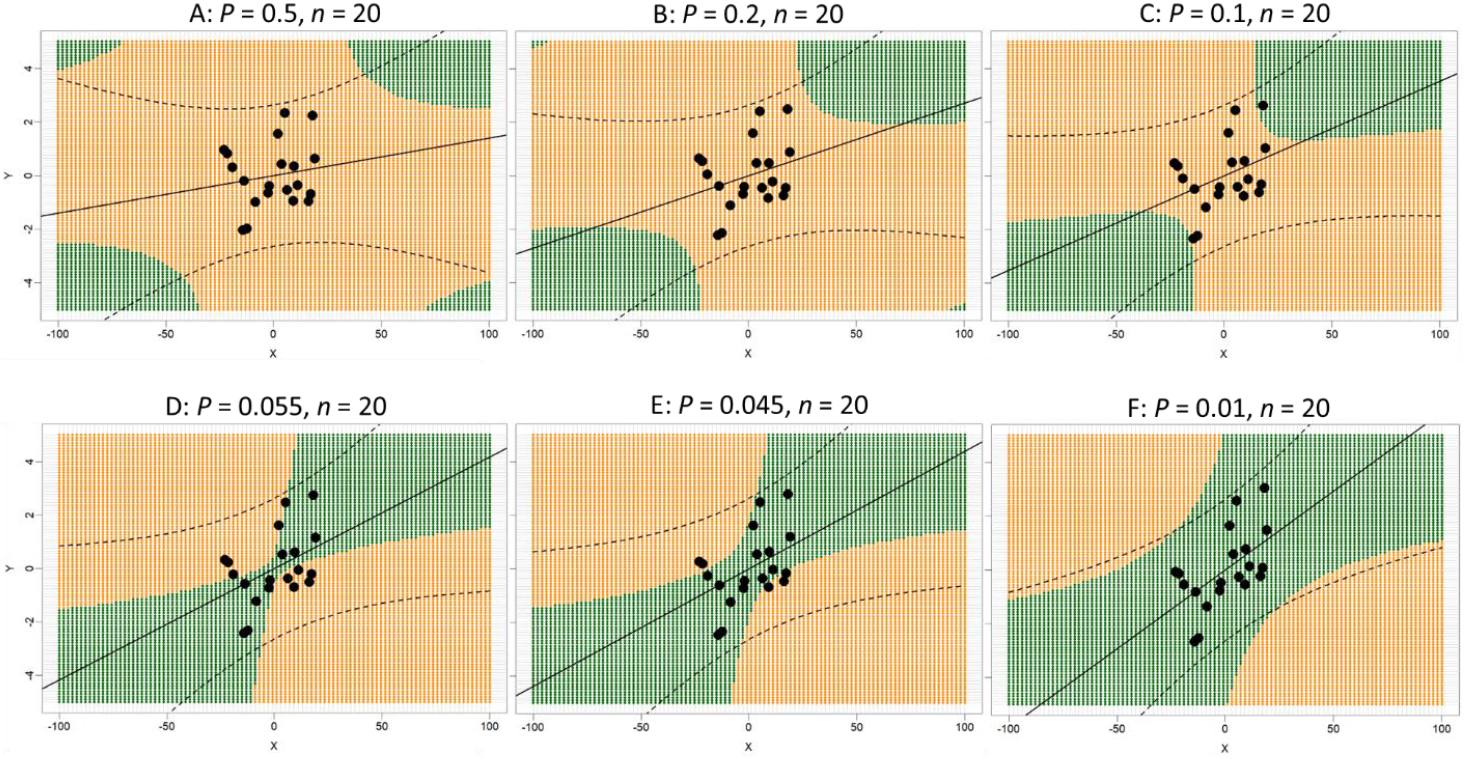
Inspection of the effect of influential data points when the regression’s *p*-value is approaching the α-threshold (*n* = 20). The area of attaining significance (green area) is increasing when the *p*-value is in proximity of the α-threshold (A-D). When the regression is already significant (E-F), decreasing *p*-values result in a narrowing of the area where an influential data point might render the regression insignificant (orange). Dashed lines: 95 % prediction interval.

**Supplemental Figure 2.**
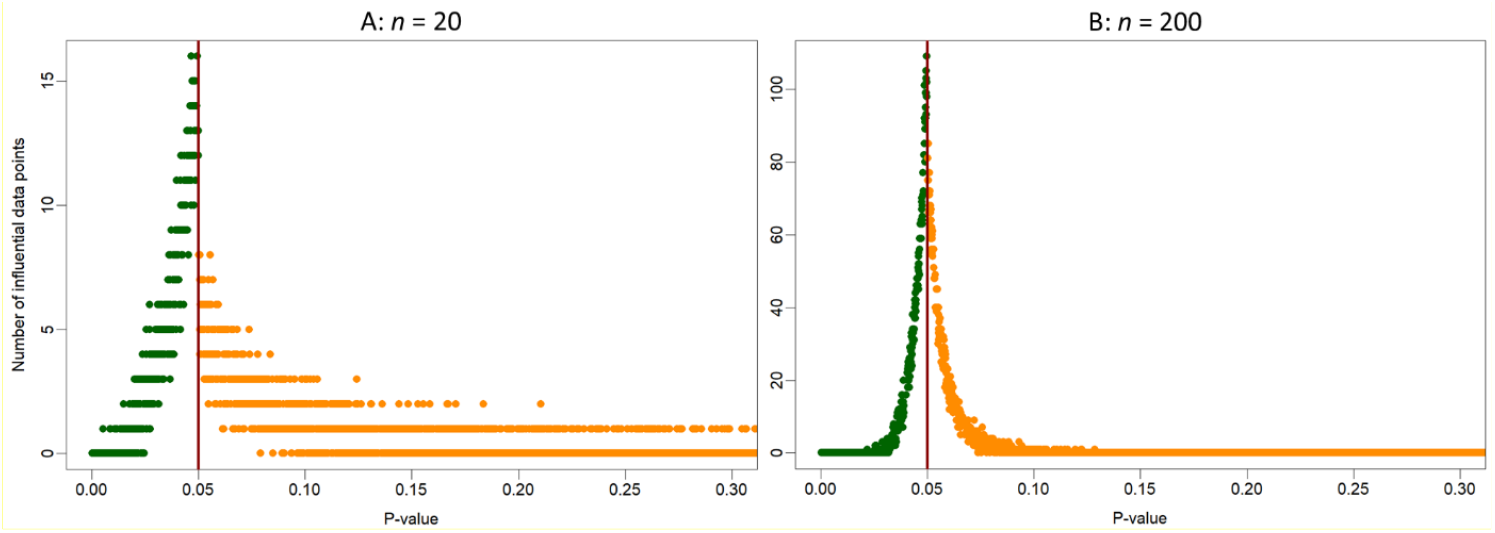
Inspection of the number of influential data points when the regression’s *p*-value is near the α-threshold (*n* = 20, 200). Note that an increasing proximity to the α-threshold (red vertical line) entails an increasing number of data points that become influential.

**Supplemental Figure 3.**
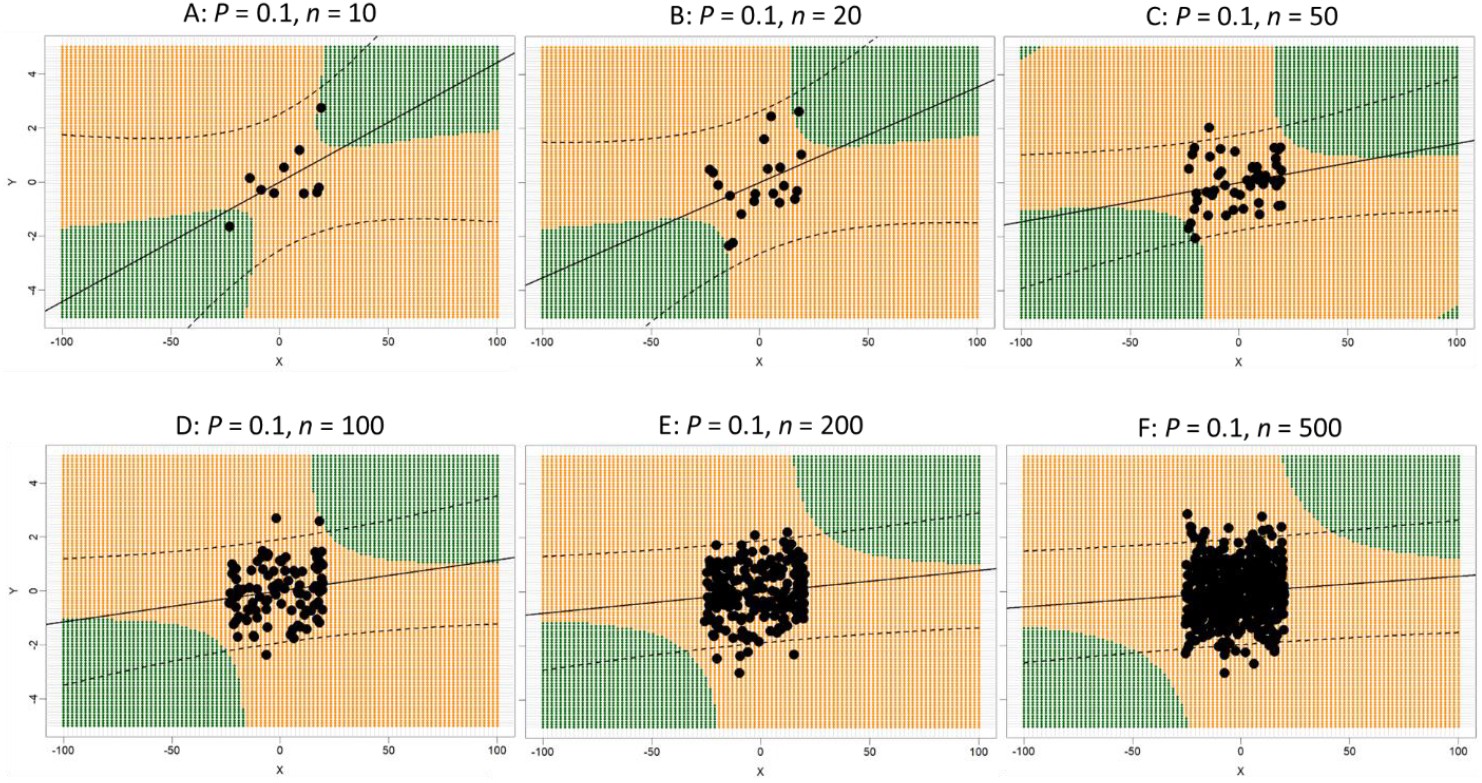
Evaluation of the effect of increasing sample sizes (*n* = 10, 20, 50, 100, 200, 500) on the susceptibility of the model to be altered with respect to the significance statement through an influential data point. Note that with increasing sample size, the area for influential data is migrating further away from the data cloud. Dashed lines: 95 % prediction interval.

## Acknowledgements

This work was funded by the “BMBF-Innovationsinitiative für die neuen Länder-Unternehmen-Region-Wachstumskern-Potenzial, miRMAK, 30WKP55B” as well as in part by the Gesundheitscampus Brandenburg “digilog: Digitale und analoge Begleiter für eine alternde Bevölkerung” initiative of the Brandenburgian Ministry of Science, Research and Culture (MWFK) to SR.

## Code availability statement

Open-source (GPL >= 2) *R* code for the “reverseR” package v0.2 is available at CRAN under https://cran.r-project.org/web/packages/reverseR/index.html.

